# Neuronal representation of a working memory-based decision strategy in the motor and prefrontal cortico-basal ganglia loops

**DOI:** 10.1101/2022.09.07.506894

**Authors:** Tomohiko Yoshizawa, Makoto Ito, Kenji Doya

**Affiliations:** Oral Physiology, Department of Oral Functional Science, Faculty of Dental Medicine and Graduate School of Dental Medicine, Hokkaido University, Kita 13, Nishi 7, Kita-ku, Sapporo, Hokkaido, Japan, 060-8586; Neural Computation Unit, Okinawa Institute of Science and Technology Graduate University, 1919-1 Tancha, Onna-son, Kunigami-gun, Okinawa, Japan, 904-0412; LiNKX, Inc. Kazama Building 5F 2-19-5 Nishi-Shimbashi, Minato-ku, Tokyo, Japan, 105-0003

## Abstract

While animal and human decision strategies are typically explained by model-free and model-based reinforcement learning, their choice sequences often follow simple procedures based on working memory of past actions and rewards. Here we address how working memory-based choice strategies, such as win-stay-lose-switch (WSLS), are represented in the prefrontal and motor cortico-basal ganglia loops by simultaneous recording of neuronal activities in the dorsomedial striatum (DMS), the dorsolateral striatum (DLS), the medial prefrontal cortex (mPFC), and the primary motor cortex (M1). In order to compare neuronal representations when rats employ working memory-based strategies, we developed a new task paradigm, a continuous/intermittent choice task, consisting of choice and no-choice trials. While the continuous condition (CC) consisted of only choice trials, in the intermittent condition (IC), a no-choice trial was inserted after each choice trial to disrupt working memory of the previous choice and reward. Behaviors in CC showed high proportions of win-stay and lose-switch choices, which could be regarded as “a noisy WSLS strategy.” Poisson regression of neural spikes revealed encoding specifically in CC of the previous action and reward before action choice and prospective coding of WSLS action during action execution. A striking finding was that the DLS and M1 in the motor cortico-basal ganglia loop carry substantial WM information about previous choices, rewards, and their interactions, in addition to current action coding.

**Significance Statement:** Working memory-based decision strategies, such as win-stay-lose-switch (WSLS), are widely observed in humans and animals. To address neuronal bases of these strategies, we recorded neuronal activities of rat prefrontal and motor cortico-basal ganglia loops during continuous/intermittent choice tasks. The rat choice strategy was a noisy WSLS in the continuous choice condition, whereas non-WSLS was selected in the intermittent choice condition. In the continuous choice condition, the primary motor cortex and the dorsolateral striatum in the motor loop more strongly conveyed information about previous choices, rewards, and their interactions than the medial prefrontal cortex and the dorsomedial striatum in the prefrontal loop. These results demonstrate that the motor cortico-basal ganglia loop contributes to working memory-based decision strategies.

## Introduction

Human and animal decision making processes can be modeled by reinforcement learning (RL) theory, in which agents update the expected reward for each choice (Sutton and Barto, 1998). However, learning can be more dynamic and hypothesis driven. Under the assumption that either one of two choices is rewarding and the other is not rewarding, “Win-Stay-Lose-Shift” or “Win-Stay-Lose-Switch” (WSLS) is an optimal strategy. WSLS can be implemented with a very high learning rate in model-free reinforcement learning (Barraclough et al., 2004; Ito and Doya, 2009), or using working-memory (WM) of previous actions and rewards (Kesner and Churchwell, 2011; Nolen-Hoeksema et al., 2014). Patients with psychiatric disorders or developmental disabilities frequently show abnormal patterns of WSLS (Shurman et al., 2005; Waltz et al., 2007; Waltz and Gold, 2007; Prentice et al., 2008; Waltz et al., 2011; Schlagenhauf et al., 2014), which may be due to disorders of WM (Barch and Ceaser, 2012).

Previous studies tested how availability of WM affected choice strategies by increasing the number of visual stimuli to remember (Collins and Frank, 2012; Collins et al., 2014), requiring an additional memory task in parallel (Otto et al., 2013a), or resulting in acute stress (Otto et al., 2013b). These studies involved humans and long inter-trial intervals in rodents (Iigaya et al., 2018), suggested that choice strategies with intact WM were close to WSLS, whereas strategies under WM disruption became similar to behavior under standard RL.

While the basal ganglia play a major role in model-free reinforcement learning (Samejima et al., 2005; Ito and Doya, 2009, 2015a; Yoshizawa et al., 2018), the neural basis of WM-based decision making is still unclear. Here we developed a choice task for rats, in which WM availability was manipulated by inserting a no-choice trial between choice trials. This task addressed how working-memory-based choice strategies, such as WSLS, are represented in the prefrontal and motor cortico-basal ganglia loops by simultaneous recording of neuronal activities in the dorsomedial striatum (DMS) and the medial prefrontal cortex (mPFC). These structures form a corticostriatal loop related to goal-directed behaviors (Voorn et al., 2004; Yin et al., 2004, 2005a; Yin et al., 2005b; Yin et al., 2006; Balleine et al., 2007; Balleine and O’Doherty, 2010). The dorsolateral striatum (DLS) and the primary motor cortex (M1) form a corticostriatal loop related to motor actions. While previous studies suggested a major role of the PFC in working memory, the present results suggest the contribution of the motor loop to WM-based decision making.

## Materials and Methods

### Subjects

Male Long-Evans rats (n = 6; 260-310 g body weight; 16-37 weeks old at the first recording session) were housed individually under a light/dark cycle (lights on at 7:00, off at 19:00). Experiments were performed during the light phase. Food was provided after training and recording sessions so that body weights decreased no lower than 90% of initial levels. Water was supplied *ad libitum*. The Okinawa Institute of Science and Technology Animal Research Committee approved the study.

### Apparatus

All training and recording procedures were conducted in a 40 × 40 × 50 cm experimental chamber placed in a sound-attenuating box (O’Hara & Co.). The chamber was equipped with three nose-poke holes on one wall and a pellet dish on the opposite wall (Figure 1A). Each nose-poke hole was equipped with an infrared (IR) sensor to detect nose entry, and the pellet dish was equipped with an infrared sensor to detect the presence of a sucrose pellet (25 mg) delivered by a pellet dispenser. The chamber top was open to allow connections between electrodes mounted on the rat’s head and an amplifier. House lights, two video cameras, two IR LED lights and a speaker were placed above the chamber. A computer program written with LabVIEW (National Instruments, Austin, Texas, USA) was used to control the speaker and the dispenser and to monitor states of the IR sensors.

**Figure 1.**
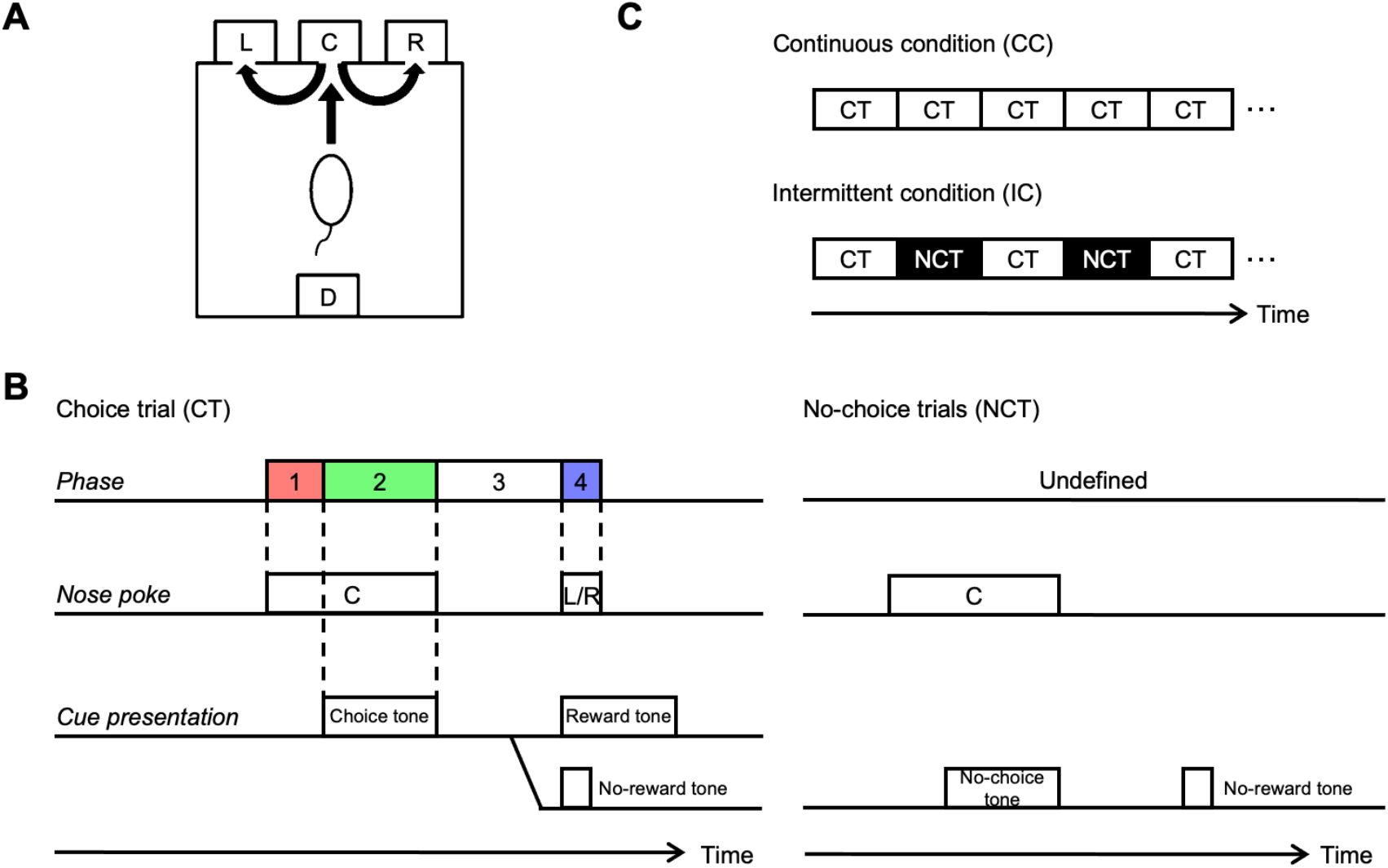
Apparatus and behavioral task. A. The experimental chamber was equipped with three holes for nose poking and a pellet dish. B. Time sequences of choice task. The behavioral task consisted of choice and no-choice trials. C. The behavioral task comprised two conditions. Choice trials were repeatedly represented in the continuous condition (CC). Choice and no-choice trials were presented alternatively in the intermittent condition (IC).

### Behavioral task

Animais were trained to perform a choice trial and a no-choice trial using nose-poke responses. In either task, each trial began with a tone presentation (start tone: 3000 Hz, 1000 ms). When the rat performed a nose-poke in the center hole for 500 - 1000 ms, one of two cue tones (choice tone: white noise, 1000 - 1500 ms; no-choice tone: 900 Hz; 1000 - 1500 ms) was presented (Figure 1B).

After onset of the choice tone (choice trials), the rat was required to perform a nose-poke in either the left or right hole within 2 s after exiting the center hole. If the rat exited the center hole before the offset of the choice tone, the choice tone was stopped. When the rat nose-poked either the left or right hole, either a reward tone (500 Hz, 1000 ms) or a no-reward tone (500 Hz, 250 ms) was presented probabilistically, depending on the selected action. The reward tone was followed by delivery of a sucrose pellet (25 mg) in the food dish. If the rat did not perform a nose-poke in either the left or right hole within 2 s, the trial was ended as an error trial after presentation of an error tone (9500 Hz, 1000 ms).

For the no-choice tone (no-choice trials), the rat was required not to perform left nor right nose pokes during 2 s after the exit from the center hole. Then, the trial was correctly finished by presentation of the no-reward tone. In this no-choice trial, the rat could not obtain any pellets, but if the rat could not perform this trial correctly, that is, if the rat incorrectly performed left or right nose-poke despite the no-choice tone, the trial was ended as an error trial after the error tone presentation, and the no-choice trial was repeated again in the next trial.

We designed the continuous condition (CC) to consist only of choice trials, and the intermittent condition (IC) had a no-choice trial inserted after every choice trial (Figure 1C). Because it was hard for the rats to continue performing in IC, we had to limit the number of choice trials to 20 in one sequential IC condition. A block is defined as a sequence of trials under the same reward probabilities of either (Left, Right) = (75%, 25%) or (25%, 75%). The first three blocks in each session were CC and the subsequent two blocks were IC. Reward probabilities of the first block were randomly selected from these two patterns for every recording session, and were switched for every subsequent block.

To adjust block-change conditions in CC and IC, the first (CC) and the third (CC) blocks were terminated when the choice frequency of the rats in the last 10 choice trials reached 80% optimal. The second (CC), and, fourth and fifth (IC) blocks were ended when 10 choice trials had been conducted. In this setting, the first 20 choice trials in the second and the third CC blocks, and in the fourth and fifth IC blocks should be comparable; starting from 80% biased choice and switching reward probabilities after 10 choice trials. This set of five blocks was repeated about six times in a one-day recording session.

### Surgery

After rats mastered the choice tasks, they were anesthetized with pentobarbital sodium (50 mg/kg, *i*.*p*.) and placed in a stereotaxic frame. The skull was exposed and holes were drilled in the skull over the recording site. Four drivable electrode bundles were implanted and fixed in the DMS in the left hemisphere (1.0 mm posterior, 1.6 mm lateral from bregma, 3.7 mm ventral from the brain surface), the DLS in the right hemisphere (1.0 mm anterior, 3.5 mm lateral from bregma, 3.3 mm ventral from the brain surface), the PL in the left hemisphere (3.2 mm anterior, 0.7 mm lateral from bregma, 2.0 mm ventral from the brain surface), and the M1 in the right hemisphere (1.0 mm anterior, 2.6 mm lateral from bregma, 0.4 mm ventral from the brain surface) using pink dental cement with confirmed effects on the brain (Yoshizawa and Funahashi, 2020).

An electrode bundle was composed of eight Formvar-insulated, 25 µm bare diameter nichrome wires (A-M Systems) and was inserted into a stainless-steel guide cannula (0.3 mm outer diameter; Unique Medical). Tips of the microwires were cut with sharp surgical scissors so that ∼ 1.5 mm of each tip protruded from the cannula. Each tip was electroplated with gold to obtain an impedance of 100-200 kΩ at 1 kHz. Electrode bundles were advanced by 125 µm per recording day to acquire activity from new neurons.

### Electrophysiological recordings

Recordings were made while rats performed choice tasks. Neuronal signals were passed through a head amplifier at the head stage and then fed into the main amplifier through a shielded cable. Signals passed through a band pass filter (50 ∼ 3000 Hz) to a data acquisition system (Power1401; CED), by which all waveforms that exceeded an amplitude threshold were time-stamped and saved at a sampling rate of 20 kHz. The threshold amplitude for each channel was adjusted so that action potential-like waveforms were not missed while minimizing noise. After a recording session, the following off-line spike sorting was performed using a template-matching algorithm and principal component analysis with Spike2 (Spike2; CED): Recorded waveforms were classified into several groups based on their shapes, and a template waveform for each group was computed by averaging. Groups of waveforms that generated templates that appeared to be action potentials were accepted, and others were discarded. Then, to test whether accepted waveforms were recorded from multiple neurons, principal component analysis was applied to the waveforms. Clusters in principal component space were detected by fitting a mixture Gaussian model, and each cluster was identified as signals from a single neuron. This procedure was applied to each 50-min data segment. If stable results were not obtained, data were discarded. Then, gathered spike data were refined by omitting data from neurons that satisfied at least one of the three following conditions:

1. The amplitude of waveforms was < 7 x the SD of background noise.
2. The firing rate calculated by perievent time histograms (PETHs) (from -4.0 s to 4.0 s with 100-ms time bin based on the onset of cue tone, exit from the center hole, or entrance into the left or right hole) was < 1.0 Hz for all time bins of all PSTHs.
3. The estimated recording site was considered outside the target.

Furthermore, considering the possibility that the same neuron was recorded from different electrodes in the same bundle, we calculated cross-correlation histograms with 1-ms time bins for all pairs of neurons that were recorded from different electrodes in the same bundle. If the frequency at 0 ms was 10 x larger than the mean frequency (from - 200 ms to 200 ms, except the time bin at 0 ms) and their PETHs had similar shapes, either one of the pair was removed from the database.

### Histology

After all experiments were completed, rats were anesthetized as described in the surgery section, and a 10µA positive current was passed for 30 s through one or two recording electrodes of each bundle to mark their final recording positions. Rats were perfused with 10% formalin containing 3% potassium hexacyanoferrate (II), and brains were carefully removed so that the microwires did not cause tissue damage. Sections were cut at 60µm on an electrofreeze microtome and stained with cresyl violet. Final positions of electrode bundles were confirmed using dots of Prussian blue. The position of each recorded neuron was estimated from the final position and the distance that the bundle of electrodes moved. If the position was outside the DMS, DLS, PL or M1, recorded data were discarded.

### Logistic Regression Analysis for behavioral data

We performed logistic regression analysis to examine the influence of past actions and outcomes on the next choice using the regression model:

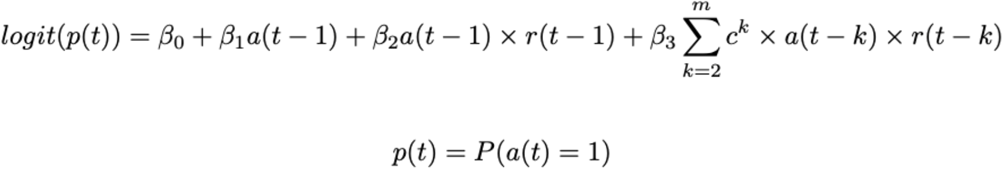

where *β*_i_ is the regression coefficient for each variable (regressor), *a(t)* ∈[1:Left, -1: Right] is the selected action, and *r(t)* ∈[1: Rewarded, -1: non-rewarded] is the reward outcome. The parameter *c* (0≤*c*≤1) specifies the decay rate of past actions and rewards. For each setting of *c*, regression coefficients *β*_i_ are derived by the “fitglm” function of MATLAB and the optimal *c* was selected between 0 to 1 with a line search. The optimal *m* was determined by comparing adjusted R^2^ with *c* set at the optimal value. The adjusted R^2^ became maximal with *m*=9.

### Poisson Regression Analysis for neuronal data

We performed two Poisson regression analyses to examine what kinds of variables were encoded in neuronal spikes. The first Poisson regression analysis considered a Poisson model in which the number of spikes at a certain phase is sampled from a Poisson distribution with the expected number of spikes at trial *t, µ* (*t*)

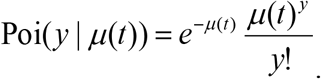

*µ* (*t*)is represented by

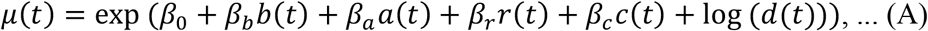

Where *²*_i_ is the regression coefficient for each explanatory variable (regressor). *b*(*t*) is the monotonically increasing factor, namely, *b*(*t*) **=** *t*, which is inserted to capture task-event-independent monotonic increases or decreases in firing pattern. *a*(*t*) *∈* [1: Contralateral, - 1: Ipsilateral] is the selected action, *r*(*t*) *∈* [1: Rewarded, 0: non-rewarded] is reward availability, and *c(t) ∈* [1: CC, 0: IC] is the task condition, *d(t)* is the time duration of a phase.

Optimal regression coefficients are determined so that the objective function, the log likelihood for all trials, is maximized.

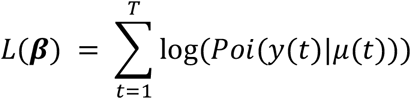

Here, **β** represents a set of coefficients. For this calculation, a function in MATLAB Statistics and Machine Learning Toolbox “fitglm(X, y, ‘Distribution’, ‘poisson’)” was used.

Next, we found the minimal necessary regressors in (A) to predict *µ* (*t*). Then, we used the Bayesian information criterion (BIC),

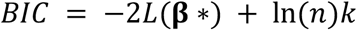

where **β\*** is the optimized beta that maximizes the log likelihood *L. k* is the number of parameters, and *n* is the trial number in a session. BIC can be regarded as a fitting measure taking into account the penalty for the number of parameters (the number of *β)* in the model.

Better models have smaller BICs. Because the full regression model includes five regressors (including the constant variable for 0), we can consider 2^5^ models for all combinations. We calculated the BIC for all possible models, and then we selected a set of regressors that showed the smallest BIC. Then, we tested the statistical significance of each regression coefficient in the selected model using the regular Poisson regression analysis. If*p* < 0.01, the corresponding variable was regarded as being coded in the firing rate. This variable selection was conducted independently for each neuron and for each time bin.

Then, we used the following full model to find neuron coding actions, rewards, interactions between actions and rewards, or prospective action using a WSLS strategy.

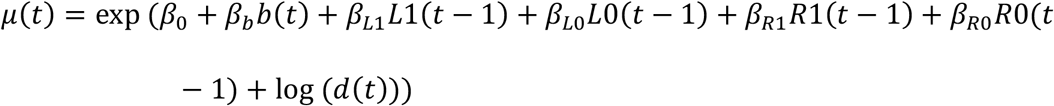

where L1(*t*) is the variable taking 1 if the rat selected left and rewarded in the trial *t*. Otherwise it takes 0. L0(*t*), R1(*t*), and R0(*t*) are the variables taking 1 only for L0, R1, or R0, respectively.

Strictly speaking, this final variable R0(t) is unnecessary, because R0(t) can be represented by 1 - L1(t) - L0(t) - R(t). Therefore, we were unable to find a unique solution of the full model that minimized likelihood. So, we considered 2^6^-1 combinations of coefficients without the full model, and found the model with the smallest BIC.

According to the variables included in the best model and the signs of these coefficients, we classified neurons into 5 types: neurons coding action in the previous choice trial, neurons coding reward in the previous choice trial, neurons coding action-reward-interaction (A×R) in the previous choice trial, prospective-action-coding, and the non-coding neurons (Table 1).

**Table 1.** Classification of information-coding neurons Pros Action: Prospective action using a WSLS strategy

|  | L1 | L0 | R1 | R0 |
| --- | --- | --- | --- | --- |
| Action | +/- | +/- |  |  |
| Action |  |  | +/- | +/- |
| Reward | +/- |  | +/- |  |
| Reward |  | +/- |  | +/- |
| A×R | + or - |  |  |  |
| A×R |  | + or - |  |  |
| A×R |  |  | + or - |  |
| A×R |  |  |  | + or - |
| Pros Action |  | +/- | +/- |  |
| Pros Action | +/- |  |  | +/- |

If the activity codes action and not reward, coefficients of the model for L1 and L0, or for R1 and R0 should have the same signs. If the activity codes reward, but not action, coefficients of L1 and R1, or L0 and R0, should be necessary and should have the same signs. If the activity codes an interaction between action and reward (AR), one coefficient among L1, L0, R1, and R0, should be necessary.

If activity codes a prospective action by a WSLS strategy (Pros Action), the coefficient of L1 and R0 (both predict action R in the next trial), or L0 and R1 (both predict action L in the next trial) should be necessary. Because multiple tests changed the ratio of false positive errors, thresholds indicating a significant proportion of these information-coding neurons were calculated so that the ratio of false positives becomes 0.05 (Binomial tests). To compare proportions of these information-coding neurons between CC and IC, data in CC and IC were analyzed separately with this Poisson regression analysis.

### Experimental design and statistical analyses

The presented analyses include 48,554 behavioral and neural trials (after the task learning was completed) recorded over a total of 78 sessions in six rats. The minimum and maximum number of trials per session were 436 and 1105, respectively.

We used appropriate statistical tests when applicable, i.e., paired or unpaired *t-* test, Mann-Whitney *U*-test, Wilcoxon signed rank test, Binominal test and Chi-squared tests with or without Bonferroni’s multiple comparison tests. Differences were considered statistically significant when p < 0.05. See “Results” for details.

## Results

To investigate whether and how the availability of working memory (WM) influences the choice strategy of rats, we designed a choice task composed of choice trials and no-choice trials. In the continuous condition (CC), choice trials were repeated and in the intermittent condition (IC), WM was disturbed by inserting a no-choice trial between each pair of choice trials (Figure 1). In each choice trial, after the choice-tone presentation, the rat performed a nose-poke in ether the left or right hole, which caused delivery of a sucrose pellet with a probability depending on the choice (Left, Right) = (75%, 25%) or (25%, 75%). Reward probabilities were changed in blocks without any explicit cue so that rats had to integrate past experiences of choices and rewards to find a better action. In no-choice trials, rats were required not to nose-poke in either the left or right hole. If the rat failed to perform the no-choice trial, the no-choice trial was repeated (see Materials and Methods for more detail).

An experimental session consisted of the first 3 blocks in CC and the fourth and fifth blocks in IC (Figure 2A). For the analysis, we used behavioral and neuronal data in 20 choice trials in the second and third blocks as a “CC sequence”, and 20 choice trials in the fourth and fifth blocks as an “IC sequence” (see Materials and Methods for more detail). In one experimental day, this sequence of 5 blocks was repeated 4-6 times (usually, 6 times; mean = 5.9 times).

**Figure 2.**
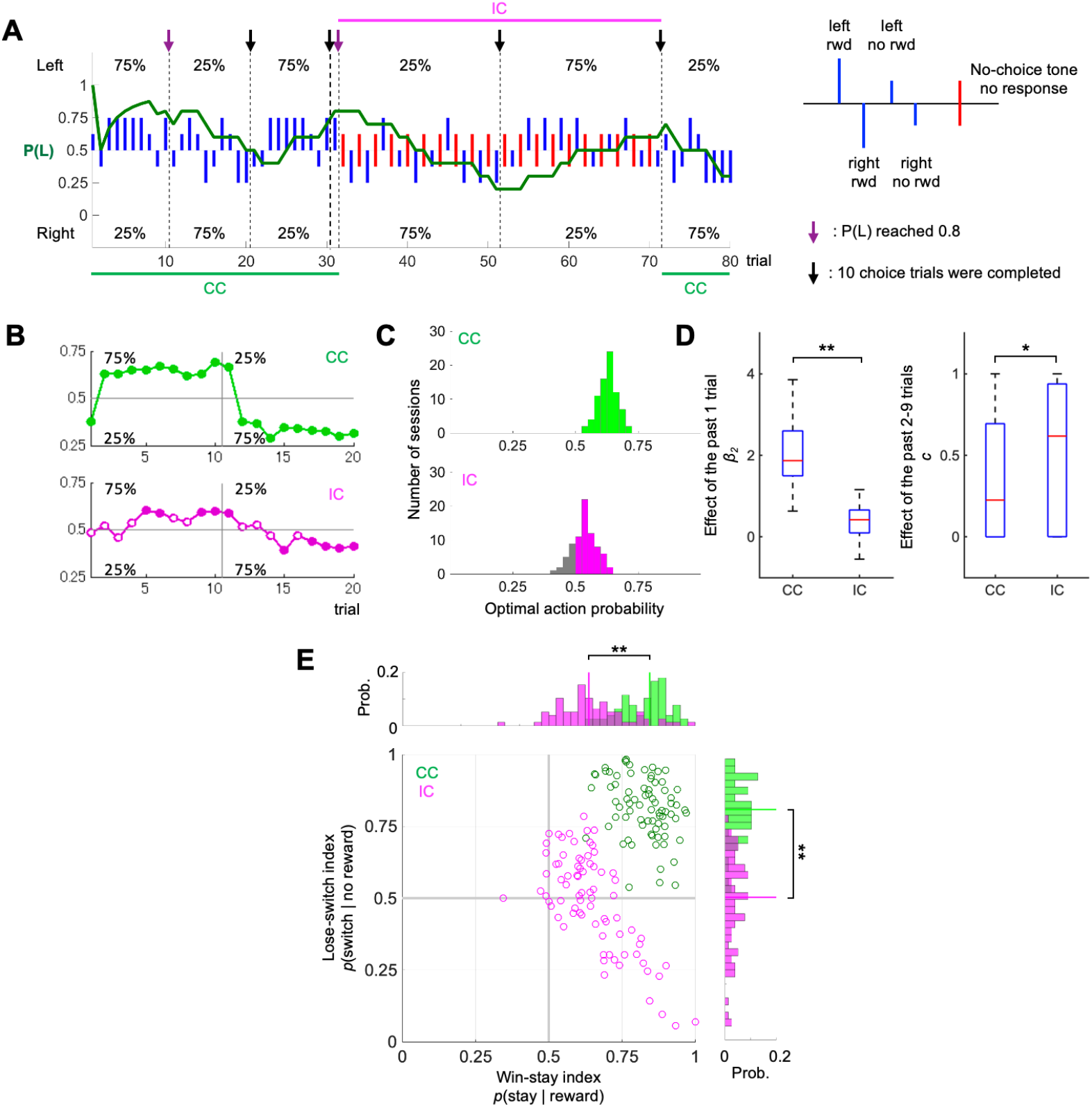
Behavioral results. A. A representative example of rat performance. Blue vertical lines indicate individual choices in choice trials. Red vertical lines indicate no-choice trials. Long lines and short lines represent rewarded and non-rewarded trials, respectively. The green trace in the middle indicates the probability of a left choice in choice trials (average of the last 10 choice trials). B. Averaged learning curves in one sequence of CC (upper) and IC (lower). A sequence consisted of 20 trials, and after 10 trials, reward probabilities were reversed. The vertical axis indicates the frequency of the action associated with the higher reward probability in the first 10 blocks. Filled circles and open circles show that the action frequency was significantly differ from 0.5 (p < 0.05; Mann-Whitney U test). C.Distributions of the optimal action probability in one session of CC (upper) and IC (lower). The optimal action probability is the frequency of selecting the action associated with the larger reward probability in one session. Medians of both distributions are significantly different from 0.5 (p < 0.01 for CC and IC; Mann-Whitney U test). D. Effects of interaction between past actions and outcomes on the subsequent action. The subsequent action was regressed by action in the previous trial and interaction of actions and outcomes in the past 9 trials. **: p<0.01, *: p<0.05, Wilcoxon signed rank test. E. Win-stay lose-switch (WSLS) indexes. The horizontal axis represents a win-stay index, the frequency that rats selected the same action after the rewarded trial. The vertical axis represents a lose-switch index, the frequency that rats switch the action after the no- reward trial. WSLS indices of CC sessions are plotted with green dots, while indices of IC sessions are shown with pink dots. Vertical lines in histograms indicate the medians of win-stay or lose-switch probabilities in CC and IC. **: p<0.01, Mann-Whitney U test..

### Behavioral performance

We trained six Long-Evans rats to perform the choice task (Figure 1, see Materials and Methods) and conducted 78 sessions consisting of 461 CC sequences and 461 IC sequences. In both conditions, rats learned to choose the optimal side with a larger reward probability (75%), based upon their experiences with choice and reward (Figure 2A). When the more rewarding side switched to the opposite side, after ten choice trials, their choices also shifted to the opposite side.

We compared choice sequences in CC and IC in response to the change of reward probabilities (Figure 2B). In CC, the choice probability switched to the other option immediately after the change of reward probabilities. In IC, the choice probability gradually shifted to the opposite side and reached a significantly different level from the chance after an average of 5 trials from the block change.

The optimal action probability (the ratio by which the rats selected the action associated with the higher reward probability) in each session was distributed between 0.5 and 0.75 with a median value of 0.632 in CC, and between 0.4 and 0.65 with a median of 0.538 in IC (Figure 2C). In both conditions, the median was significantly greater than the chance level (p = 1.6e^-14^ in CC and p = 1.1e^-7^ in IC, Wilcoxon signed-rank test), confirming that choice behavior in both CC and IC adapted to reward-probability changes.

To investigate choice strategies in more detail, using logistic regression, we examined the influence of past actions and outcomes on the subsequent choice (see Materials and Methods). The action choice *a*(*t*) was regressed from the last action, *a*(*t*-1), the interaction between the last action and the outcome *a*(*t* — 1) x *r*(*t* — 1), and exponentially decaying memory of past interactions 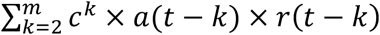, where *a*(*t*) ∈ [1: Left, -1: Right] is the selected action, *r*(*t*) ∈ [1: rewarded, -1: non-rewarded], and 0 ≤*c* ≤1 is the decay constant. The interaction of the action and the outcome of the last trial affected the subsequent action more strongly in CC than in IC (Figure 2D; median of coefficient for a(t-1)*r(t-1) β2: CC; 1.87, IC; 0.42, p = 3.3e-12, Wilcoxon signed rank test). The effect of the previous action-reward relationship decayed more rapidly in CC (Median of the decay constant *c:* CC; 0.23, IC; 0.62, p = 0.048 with no significant difference in the coefficient β3: CC; 0.13, IC: 0.19, p = 0.49). These results indicate that rats recognized the side with a larger reward probability and changed their action selection in both CC and IC, while insertion of no-choice trials in IC made learning slower.

Next, in order to test the hypothesis that the choice strategy is closer to WSLS in CC than in IC, we calculated WSLS indices, composed of the win-stay ratio P(stay| reward), the frequency that the rats chose the same action after the rewarded choice trial, and the lose-switch ratio P(switch| no-reward), the frequency that the rats switch the action after no-reward trials. WSLS indices calculated for each session are plotted two-dimensionally, P(stay| reward) vs. P(switch| no-reward) (Figure 2E). WSLS indices of CC sessions were concentrated near the upper right corner, while those of IC sessions were widely distributed from the center to the lower right corner. These two distributions had little overlap. Ratios of win-stay P(stay| reward) and lose-switch P(switch| no-reward) were significantly larger in CC than in IC (win-stay: 0.85 vs 0.64, p = 1.2e-16, Mann-Whitney *U* test; lose-switch: 0.82 vs 0.51, p = 2.6e-23). Since behaviors in CC showed high P(stay| win) and P(switch| lose), these could be regarded as “a noisy WSLS strategy” (cf. the regular WSLS is deterministic).

### Neuronal activities of the prefrontal and motor cortico-basal ganglia loops

We recorded neuronal activity in DMS, DLS, mPFC, and M1 of rats performing the choice task. Each rat was implanted with four bundles of eight microwires. Bundles were advanced by 125 µm each recording day so that data from new neurons were acquired for 4-6 sessions consisting of IC and CC sequences. After all experiments were completed, locations of bundles were confirmed by Nissl staining (Figure 3A, B). Stable recordings were made from 320 neurons in DMS, 210 neurons in DLS, 158 neurons in mPFC, and 247 neurons in M1 from six rats. We analyzed neural activities in CC and IC conditions in four phases in a trial (see Figure 1B. 1: in the center hole before the choice tone; 2: after the choice tone before exiting the center hole; 3: during the approach to the left or right hole; 4: in the left/right hole with a reward/no reward tone).

**Figure 3.**
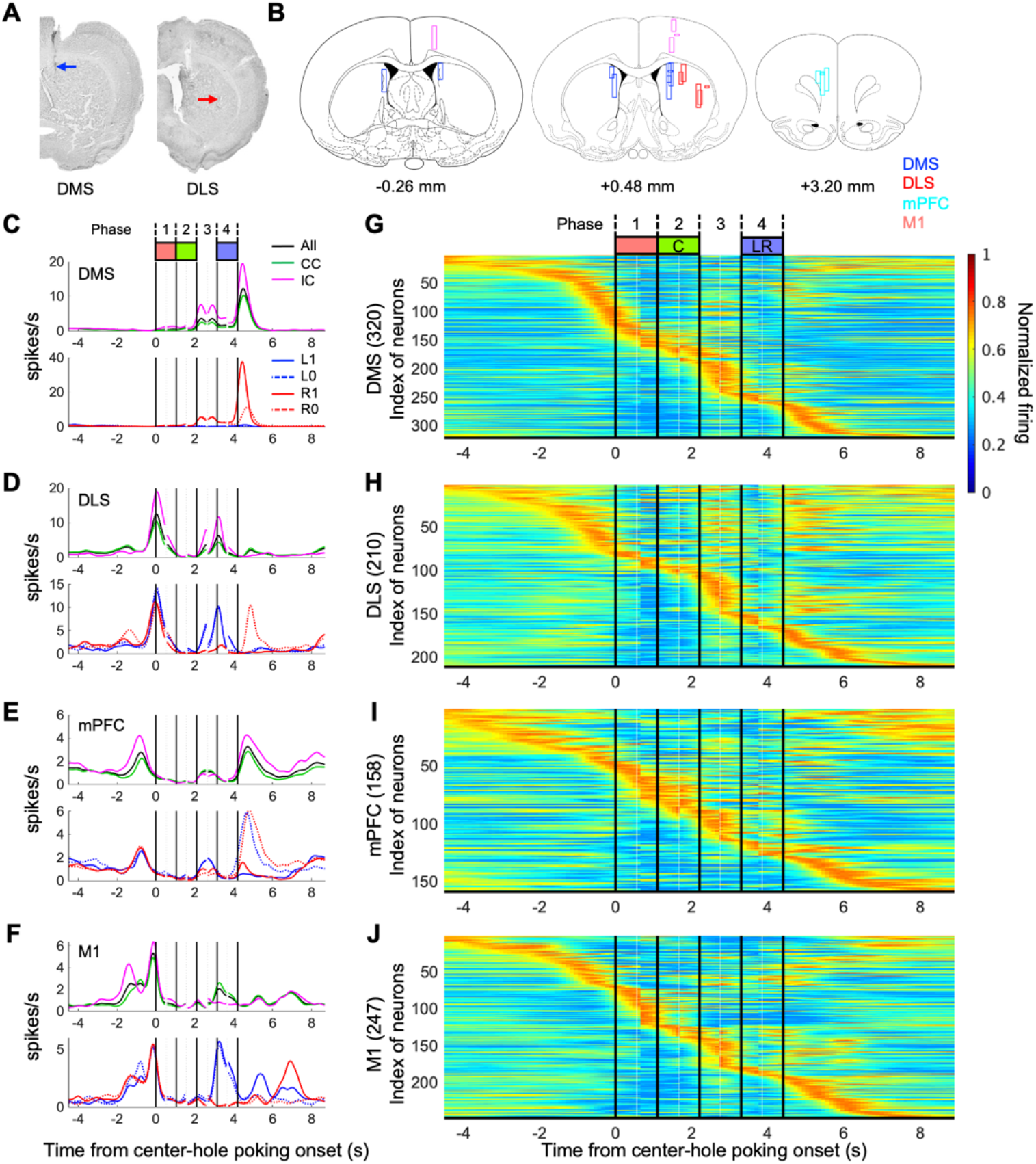
Representative activity patterns of neurons during choice trials. A. Nissl-stained coronal sections showing recording locations for the DMS (blue) and DLS (red). B. Tracks of accepted electrode bundles for all rats are indicated by rectangles. Neurons recorded from blue, red, cyan, and magenta rectangles were classified as DMS, DLS, mPFC, and M1 neurons, respectively. Each diagram represents a coronal section referenced to the bregma (Paxinos and Watson, 1998). C-F. Peri-event time histograms (PETHs) of a representative DMS neuron (C), DLS neuron (D), mPFC neuron (E), and M1 neuron (F). PETHs were calculated based on timings of five task events (onset of center-hole poking, onset of the tone, offset of center- hole poking, onset of left or right hole-poking, offset of left- or right-hole poking), and the following four task phases were defined: Phase 1: the period from the start of the center-hole poking to the onset of the cue tone. Phase 2: the choice tone presentation period. Phase 3: the action execution period between exiting the center hole and entry into the left or right hole. Phase 4: the feedback period when a reward or no-reward tone was presented after left- or right-hole poking. PETHs of all choice trials (black), and of trials in CC (green) and trials in IC (cyan) (Upper panel). PETHs of CC and IC choice trials in which left was selected and rewarded (L1), left was selected and not rewarded (L0), right was selected and rewarded (R1), or right was selected and not rewarded (R0) (Lower panel). All PETHs (50-ms bins) were smoothed with a Gaussian kernel with a 150-ms SD. C-F. Peri-event time histograms (PETHs) of a representative DMS neuron (C), DLS neuron (D), mPFC neuron (E), and Ml neuron (F). PETHs were calculated based on timings of five task events (onset of center-hole poking, onset of the tone, offset of center-hole poking, onset of left or right hole-poking, offset of left- or right-hole poking), and the following four task phases were defrned: Phase 1 : the period from the start of the center-hole poking to the onset of the eue tone. Phase 2: the choice tone presentation period. Phase 3: the action execution period between exiting the center hole and entry into the left or right hole. Phase 4: the feedback period when a reward or no-reward tone was presented after left- or right-hole poking. PETHs of all choice trials (black), and of trials in CC (green) and trials in IC (cyan) (Upper panel). PETHs of CC and IC choice trials in which left was selected and rewarded (Ll), left was selected and not rewarded (LO), right was selected and rewarded (RI), or right was selected and not rewarded (RO) (Lower panel). All PETHs (50-ms bins) were smoothed with a Gaussian kernel with a 150-ms SD. G-J. Normalized activity patterns of all recorded neurons from the DMS (G), DLS (H), mPFC (I), and M1 (J). An activity pattern for each neuron was normalized so that the maximum PETH was 1 and represented by pseudo-color (values from 0 to 1 are represented from blue to red). Indexes of neurons were sorted based on the time that the normalized PETH first surpassed 0.8.

Representative examples of spike peri-event time histograms (PETHs) with inter-trial time alignment (Ito and Doya, 2015b) are shown in Figure 3C-F. Neurons in DMS (Figure 3C) increased their activity when the rat exited from the center hole and entered the left or right hole (phase 3), and showed the largest peak after exiting the left or right hole, when the rat anticipated obtaining a pellet (black line in upper panel). PETHs for CC (green line) and IC (magenta line) differed in phase 3 and after exiting the left/right hole, showing that the activity pattern was modulated by the task condition. PETHs also showed different increased activity following a right side, especially when the choice was rewarded (R1), showing that the activity pattern was modulated by a conjunction of choice and reward.

The DLS neuron in Figure 3D increased its activity when the rat was approaching the center hole (phase 1). It showed higher activity for left than right choices in phases 3 and 4, which was higher in IC, showing context-dependent action coding.

The mPFC neuron in Figure 3E showed higher activity for no-reward than reward experiences after exiting the left/right hole, showing outcome coding.

PETHs of the M1 neuron (Figure 3F) showed higher activity when entering the left hole (phase 4), which was stronger in CC, showing context-dependent action coding.

To see the distribution of peak timings of all DMS neurons, PETHs for all DMS neurons were normalized and represented by color, with neuron indices sorted on the basis of peak activity timing (Figure 3G). Activity peaks of DMS neurons were widely distributed in different phases in a trial. PETHs of neurons in DLS, mPFC, and M1 (Figures 3H-J) also showed wide distributions of activity peaks, with greater apparent concentrations when approaching the center hole and choice holes (phase 3) in DMS and DLS, and during choice tone presentation (phase 2) in mPFC. These analyses show that neurons in the four areas show various actions, outcomes, and task condition coding at different times in the trial.

### Neuronal representation of experiences of the current choice trial

We next applied Poisson regression analyses of spike counts in each phase to quantify neuronal coding of actions, rewards, and task conditions. In these analyses, we used actions (contralateral/ipsilateral), outcomes (reward/no reward), and task conditions (CC/IC) as regressors (see Materials and Methods). Note that action coding in phase 1 and 2 represents an action command or plan before an action is performed, and that the outcome regressor means reward prediction in phase 1∼3 (Figure 4A). Average durations of each phase did not differ significantly between CC and IC (Figure 4B), which excludes the possibility that differences in neural activities were due simply to differences in motor behaviors.

**Figure 4.**
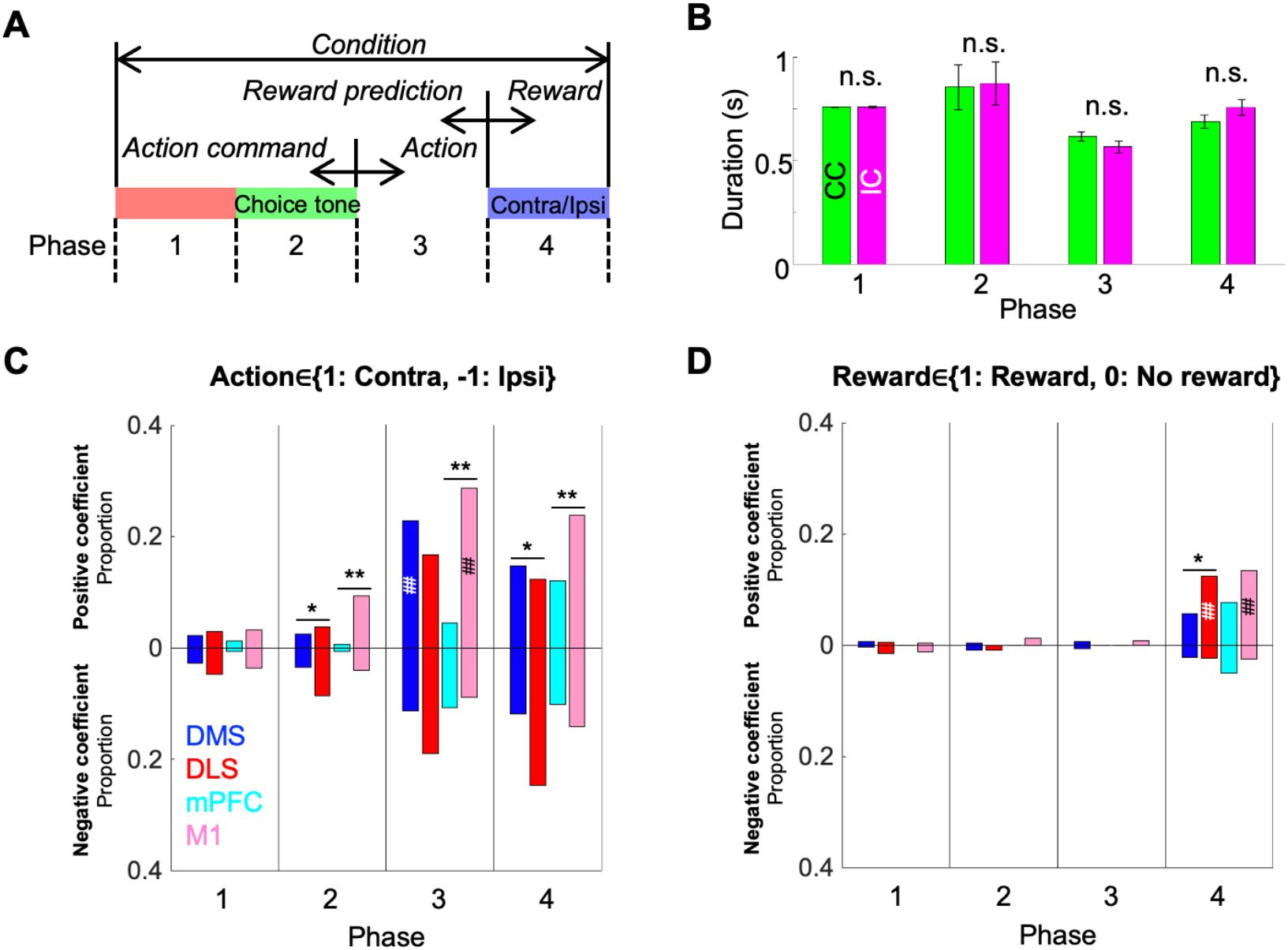
Proportions of neurons coding actions, rewards, and conditions. A. Meanings of regressors in each task phase. B. Means of time duration in each phase do not differ significantly between CC and IC. Results are means ± SEM. n.s.: p > 0.05 (Paired t-test) Results are means ± SEM. n.s.: p > 0.05 (Paired z-tcst) C. D. Proportions of neurons significantly correlated with a regressor for action (C) and reward (D) in each phase. Upward and downward bars indicate proportions of neurons with positive and negative regression coefficients, respectively. **: p < 0.01, *: p<0.05, Chi-squared test between the DMS and DLS or mPFC and M1. ##: p<0.01, #: p<0.05, Binominal test.

The proportion of neurons correlated with action was significantly larger in the motor loop than in the prefrontal loop in phases 2 and 4 (Figure 4C; phase 2: DMS: 5.9%, DLS: 12%, p = 0.037, mPFC: 1.3%, M1: 13%, p= 9.5e-05; phase 4: DMS: 27%, DLS: 37%, p = 0.039, mPFC: 22%, M1: 38%, p=0.0032, Chi-squared test with Bonferroni correction). The proportion was significantly larger in M1 than in the mPFC in phase 3 (mPFC: 15%, M1: 38%, p = 1.2e-06). A contralateral bias was clearly observed during action (phase 3) in M1, but also in DMS neurons (M1: p = 7.8e-04, DMS: p = 0.044, Binominal test with Bonferroni correction). These results indicate that the motor loop is more strongly involved in action preparation, execution, and memory than the prefrontal loop.

The proportion of neurons correlated with the outcome variable was significantly larger in the DLS than in the DMS in phase 4 (Figure 4D; DMS: 7.8%, DLS: 15%, p=0.044). Neurons in the DLS and M1 were significantly more activated by reward than no-reward experience (DLS: p = 0.018, M1: p = 0.0045). These results indicate that DLS neurons can more strongly distinguish reward-associated sensory stimulus than those in the DMS.

In all phases, the proportion of neurons correlated with the task condition was less than 10% (figure not shown), which suggests that neuronal activities did not directly encode task condition, but does not exclude the possibility that information coding of action or outcome in the previous choice trial was modulated by task conditions.

### Neuronal representation of experiences in the previous choice trial

Next, to examine how information from choice trials transferred to the subsequent choice trial with and without WM interference, we applied a second Poisson regression analysis to spike data in CC and IC separately (see Materials and Methods, Table 1). In this analysis, activity of each neuron in each phase was classified exclusively into 5 groups: neurons coding the action of the previous choice trial, neurons coding the reward of the previous choice trial, neurons coding the interaction between action and reward (A ×R) of the previous choice trial, neurons coding the action according to WSLS strategy, and non-coding neurons (see Materials and Methods).

In phase 1 (Figure 5A), upon entry into the center hole, coding of the previous reward was seen in DLS, mPFC, and M1, specifically in CC (DLS: CC; 9.0%, IC; 1.9%, p = 0.0052, mPFC: CC; 15%, IC; 1.3%, p = 2.7e-05, M1: CC; 8.5%, IC; 1.2%, p = 0.00066, Chi-squared test with Bonferroni correction). Coding of the interaction of the previous action and reward was seen in all four areas, with more DMS neurons in CC than in IC (CC; 16%, IC; 9.4%, p = 0.037).

**Figure 5.**
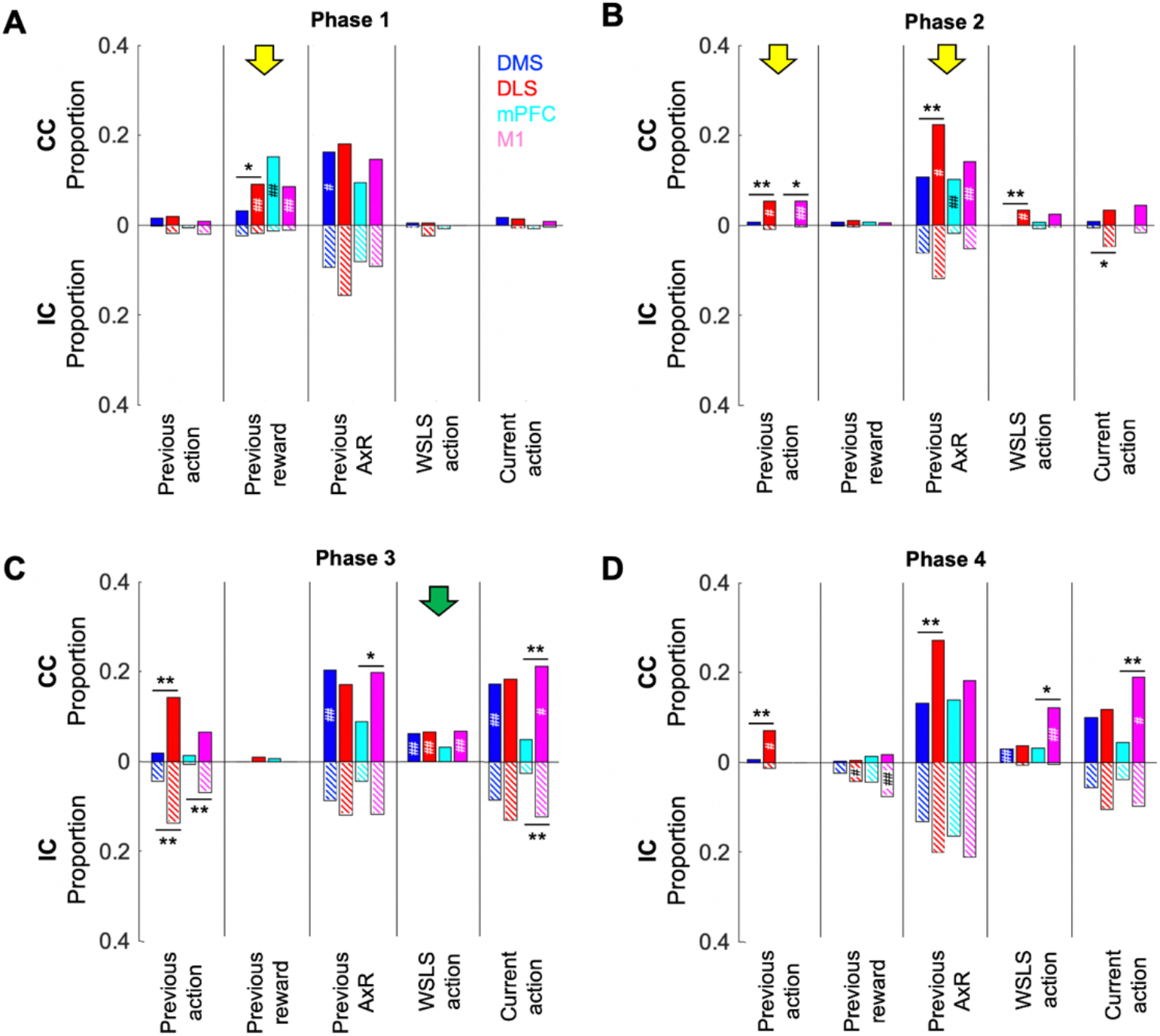
Neuronal representation of WSLS strategy in the prefrontal and motor cortico-basal ganglia loops. Proportions of neurons coding actions, rewards, and AxRs of the previous choice trial and coding WSLS and current action in phase 1 (A), 2 (B), 3 (C) and 4 (D). Filled and hatched bars indicate proportions in CC and IC, respectively. When rats employed the WSLS strategy, DLS neurons more strongly conveyed information about the previous choice trial during action preparation than DMS neurons (yellow arrows). Neuronal activities of all areas excluding the mPFC represented WSLS action during action execution (Green arrow). **: p<0.01, *: p<0.05, Chi-squared test between DMS and DLS or mPFC and M1 in each task condition. ##: p<0.01, #: p<0.05, Chi-squared test between CC and IC in each recording area.

In phase 2 (Figure 5B), while the cue tone was presented, neurons in the DLS and M1 coded the previous action selectively in CC (DLS: CC; 5.2%, IC; 0.95%, p= 0.044, M1: CC; 5.3%, IC; 0.40%, p= 0.044). More neurons in DLS, mPFC, and M1 neurons coded the interaction of the previous action and reward in CC than in IC (DLS: CC; 22%, IC; 12%, p= 0.018, mPFC: CC; 10%, IC; 1.9%, p=0.0084, M1: CC; 14%, IC; 5.3%, p= 0.0033).

In phase 3 (Figure 5C), when rats moved to left/right hole, neurons coding the WSLS action predicted from the previous action and reward were seen in the DMS, DLS, and M1 selectively in CC (DMS: CC; 6.3%, IC; 0%, p= 2.2e-05, DLS: CC; 6.7%, IC; 0%, p = 0.00057, M1: CC; 6.9%, IC; 0%, p = 0.00011). Coding of the current action was seen in all four areas, with higher proportions in CC than in IC in the DMS and M1 (DMS: CC; 17.5%, IC; 8.4%, p= 0.0026, M1: CC; 21%, IC; 12%, p= 0.023).

In phase 4 (Figure 5D), when rats entered the left/right hole and heard the reward/no-reward tone, while the proportion of M1 neurons coding the WSLS action (CC; 12%, IC; 0.40%, p= 3.0e-07) and the current actions (CC; 19%, IC; 9.7%, p= 0.013) remained higher in CC than in IC, the proportion of neurons coding the previous reward was larger in IC than in CC in the DLS and M1 (DLS: CC; 0.48%, IC; 4.3%, p= 0.042, M1: CC; 1.6%, IC; 7.7%, p= 0.0054).

Interestingly, throughout the four phases in CC, neurons coding working memory about the previous trial were more prevalent in the motor loop than in the prefrontal loop (Phase 1, previous reward: DMS vs. DLS; 3.1% vs. 9.0%, p= 0.027. Phase 2,previous action: DMS vs. DLS; 0.63% vs. 5.2%, p= 0.0063, mPFC vs. M1; 0% vs. 5.3%, p= 0.027. Phase 2, previous AxR: DMS vs. DLS; 11% vs. 22%, p= 0.0019. Phase 3, previous action: DMS vs. DLS; 1.9% vs. 14%, p= 2.2e-07. Phase 3, previous AxR: mPFC vs. M1; 8.9% vs. 20%, p= 0.023. Phase 4, previous action: DMS vs. DLS; 0.63% vs. 7.1%, p= 0.00025. Phase 4, previous AxR: DMS vs. DLS; 13% vs. 27%, p= 0.00041, Chi-squared test with Bonferroni correction), even though working memory is regarded as a major function of the prefrontal cortex.

## Discussion

Our main achievements and findings in this research are as follows.

1. We developed a choice task that manipulated WM availability for rats and showed that disturbance of WM disrupted the WSLS choice strategy (Figure 2).
2. Poisson regression of neural spikes showed that the proportions of neurons coding the current action before and after action choice were larger in the DLS than DMS, and in M1 than the mPFC (Figure 4).
3. Before action choice, the proportion of neurons coding the previous reward was larger in CC in DLS, mPFC, and M1, and the proportion coding the previous action was larger in CC in DLS and M1. During action execution in CC, neuronal activities of DMS, DLS, and M1 represented prospective action by the WSLS strategy (Figure 5).
4. Throughout the trial, working memories of previous actions, rewards, and their interactions were more prevalent in the motor loop than in the prefrontal loop (DLS than the DMS and M1 than mPFC; Figure 5).

In the present study, we showed the effect of WM availability in the choice behavior of rats, which allowed us to analyze neuronal correlates of WM-based choice strategy in the cortico-striatal circuit (Figure 2).

Recent studies in both humans and rodents showed that disruption of WM changed the strategy in sequential choice tasks (Collins and Frank, 2012; Worthy et al., 2012; Otto et al., 2013a; Otto et al., 2013b; Collins et al., 2014; Economides et al., 2015; Iigaya et al., 2018). Worthy et al. (2012) used a two-choice task and disrupted WM by requiring an additional memory task in parallel. They showed that human choice behavior with intact WM was better fitted by a WSLS model than a RL model, while choice behavior with WM load was better fitted by a RL model than the WSLS model. Collins and Frank (2012) and Collins et al. (2014) used a choice task in which a subject selected one action among three options for a given visual image. The WM load was controlled by varying the number of visual stimuli. They analyzed the choice strategy using a hybrid model combining a RL model and a WM model and suggested that the choice behavior without WM load can be explained by the WM model (RL model with the learning rate = 1), while the choice behavior with WM load can be explained by the RL model with the lower learning rate. Iigaya et al. (2018) studied mouse performance in a non-stationary, reward-driven, decision-making task and assessed WM availability based on spontaneous variations in inter-trial-intervals (ITIs). Mice showed WSLS-like choices after short ITIs, but RL-like choices after long ITIs. Optogenetic stimulation of dorsal raphe serotonin neurons boosted the learning rate only in trials after long ITIs, suggesting that serotonin neurons modulate reinforcement learning rates, and that this influence is masked by WM-based decision mechanisms.

All previous studies examining WM effects on choice strategies except Iigaya et al. (2018) were performed with human subjects, and there have been no reports comparing neuronal representations of a WM-based strategy. In this study, we recorded neurons in the DMS, DLS, mPFC, and M1 during a choice task with and without WM disturbance, and found that neurons in each area had a variety of activity patterns throughout a trial, and that patterns were modulated by selected actions, reward outcomes, and task conditions (Figure 3). Information on action command and selected action was more strongly represented in the motor loop than in the prefrontal loop (Figure 4). These properties are similar to our previous observations in the DLS and DMS neurons (Ito and Doya, 2015a).

What kind of information must be retained between trials for the WSLS strategy? There are at least two possibilities. One is to retain direct experiences with action and reward in the previous trial, such as, L1, L0, R1, and R0 (keeping past experience). Another is to compute the next action using a WSLS rule soon after reward feedback (prospective action), that is L for L1 or R0, and R for L0 or R1 and retain that until the next choice (keeping future plan).

Indeed, before action execution in CC, coding of the previous action was seen in the DLS and M1 and coding of the previous reward was seen in the DLS, mPFC, and M1 (Figure 5A and B). Combinatorial information of the previous action and reward was more dominant in CC in the DMS (phase 1) and the DLS, mPFC, and M1 (phase 2). A previous study in monkeys reported that prefrontal neurons modulated their activity according to the previous outcome and the conjunction of the previous choice and outcome (Barraclough et al., 2004). Our results showed for the first time that the motor loop (DLS and M1) retains that information more strongly than the prefrontal loop (DMS and mPFC) when a WM-based strategy was observed.

From the viewpoint of information processing, keeping a future plan is more efficient in that it requires less memory capacity. Memory of future action has been termed “prospective action” in previous studies (Kesner, 1989; Goto and Grace, 2008; Kesner and Churchwell, 2011). Although, these studies suggested that the prefrontal cortex is responsible for prospective action coding, in our study, prospective WSLS coding appeared during action execution in all recorded areas, excluding the mPFC. It is still possible that the prospective action was calculated and retained in other prefrontal areas, such as the anterior cingular cortex (ACC), located just above the mPFC. The ACC was considered a part of the prefrontal cortex in previous studies (Kesner, 1989; Goto and Grace, 2008; Kesner and Churchwell, 2011) and was thought to be involved in working memory for the motor response (Kesner and Churchwell, 2011).

In conclusion, this experiment showed that the availability of WM affects choice strategies in rats and revealed WM-related neuronal activities in DMS, DLS, mPFC, and M1. A striking finding was that DLS and M1 in the motor cortico-basal ganglia loop carry substantial WM information about previous choices, rewards, and their interactions, in addition to action coding during action execution.

## 16. Acknowledgements

This work was supported by JSPS KAKENHI Grant Numbers JP16H06563, JP23120007 to K.D., JP19K16299, JP22K15217 to T. Y. and generous research support of Okinawa Institute of Science and Technology Graduate University for the Neural Computation Unit. We thank the members of the Neural Computation Unit for helpful comments and discussion, and Steven D. Aird (https://www.sda-technical-editor.org) for thorough editing and proofreading.

## Notes

**Conflict of Interest:** The authors report no conflicts of interest.

### Competing Interest Statement

The authors have declared no competing interest.

